# S100A8/A9 modulates inflammatory collateral tissue damage during intraperitoneal origin systemic candidiasis

**DOI:** 10.1101/2020.03.30.017491

**Authors:** Madhu Shankar, Nathalie Uwamahoro, Sandra Holmberg, Maria Joanna Niemiec, Johannes Roth, Thomas Vogl, Constantin F. Urban

## Abstract

Peritonitis is a leading cause of severe sepsis in surgical intensive care units, as over 70% of patients diagnosed with peritonitis develop septic shock. A critical role of the immune system is to return to homeostasis after combating infection. S100A8/A9 (calprotectin) is an antimicrobial, pro-inflammatory protein complex often used as a biomarker for diagnosis of disease activities in many inflammatory disorders. Here we describe the role of S100A8/A9 on inflammatory collateral tissue damage (ICTD).

We performed an *in vivo Candida albicans* disseminated peritonitis mouse model using WT and S100A9-deficient mice and stimulated primary macrophages with recombinant S100A8/A9 in the presence or absence of the compound paquinimod, a specific inhibitor of S100A9. In addition, the effects on ICTD and fungal clearance were investigated. S100A9-deficient mice developed less ICTD than wildtype mice. Restoration of S100A8/A9 in S100A9 knockout mice resulted in increased ICTD and fungal clearance comparable to wildtype levels. Treatment with paquinimod abolished ICTD.

The data indicated that S100A8/A9 controls ICTD levels and host antimicrobial modulation at a systemic level during intra-abdominal candidiasis (IAC).

## Introduction

Peritonitis frequently results in severe sepsis in surgical patients, especially in the intensive care unit (ICU) (1), as more than 70% of patients often succumb to death within 72h (2). Hence, treatment options are required that extend life expectancy to allow proper treatment (3). Peritonitis is characterized by inflammation of the membrane in the abdominal cavity, the peritoneum. Peritonitis often occurs upon disruption of physical barriers or can become spontaneous in severe organ failure, causing alterations of the physiologic flora residing in the gastrointestinal tract (1,4). These alterations prompt an inflammatory response that targets the removal of contaminants from the peritoneal cavity into circulation. There, pathogens induce activation of host immune responses and the release of pro-inflammatory mediators including Interleukin-6 (IL6), macrophage inflammatory protein −1α (MIP-1α) and tumor necrosis factor –α (TNFα) to recruit phagocytes to the peritoneum (5). The attempt to restrict infection promotes abscess development through production of fibrinous exudate. Failure to confine peritonitis may lead to organ failure, coma and death (6).

The inflammatory response of the innate immune system can either have intentional (pathogen clearance) or collateral outcomes (pathological side effect, called inflammatory collateral tissue damage -ICTD) (7). The focus of most studies on detection and elimination of pathogens has neglected host tolerance to disease through the resolution of collateral outcomes of inflammation in an attempt to establish homeostasis (8).

Acute or systemic inflammatory pathological states are often associated with intra-abdominal surgery (9). Surgical intervention disrupting the natural barriers of the gastrointestinal tract leads to deep-seated microbial infection by gut colonizing organisms and frequently *Candida albicans* infection. As a result, the most common non-mucosal fungal diseases among hospitalized patients are *Candida* peritonitis, also referred to as intra-abdominal candidiasis (IAC). IAC is often challenging to diagnose and hence results in high mortality rates ranging from 25% - 60% (9).

The resolution of infection is an active process involving reprogramming of cells and modulation of mediators (10) such as the S100A8/A9 heterodimeric complex (calprotectin) also known as alarmin or DAMP (11–13). Here, we used an experimental model for IAC to study the contribution of S100A8/A9 to ICTD.

Participating as a signaling molecule that binds to Toll-like receptor-4 (TLR4) to induce pro-inflammatory chemokines and cytokines, the heterodimer is the physiologically relevant and active form that is secreted by activated, stressed or necrotic cells. The prolonged presence of S100A8/A9, however, leads to a tolerated state of the immune system as a counter mechanism or stress tolerance (12). Under high calcium conditions, such as present in the extracellular milieu and culture medium, S100A8/A9 heterodimers inactivate itself by tetramer formation, to restrict their activity locally and to avoid overwhelming immune reactions (13). Thus, S100A8 homodimers, which cannot tetramerise, are widely accepted experimental stimuli to mimic heterodimer activity.

Furthermore, S100A8/A9 binds micronutrients including zinc, manganese, and calcium, and is often deployed by leukocytes as a mechanism to either deprive microbes of the nutrients or poison the microbes in high quantities (14). Neutrophils release the heterodimer during the formation of neutrophil extracellular traps to bind or capture *C. albicans* (15). In addition to the antimicrobial benefit, S100A8/A9 is used in diagnostics to monitor neutrophil elevation in various inflammatory diseases (16). The q-compound paquinimod, an immunomodulatory compound that prevents the binding of S100A9 to TLR4, was designed to target chronic inflammatory S100A8/A9 dependent diseases (17–22). Using a murine peritonitis model with *C. albicans*, we show here that the inflammatory response failed to contain the pathogen in the peritoneum but instead led to detrimental ICTD dependent on the presence of S100A8/A9. Treatment of mice with paquinimod abrogated S100A8/A9-induced ICTD suggesting paquinimod as promising adjunct therapy option during severe IAC.

## Materials and methods

### Ethical statement

Animal experiments and isolation of cells were carried out following the recommendations in the Guide for the Care and Use of Laboratory Animals, conformed to Swedish animal protection laws and applicable guidelines (djurskyddslagen 1988:534; djurskyddsförordningen 1988:539; djurskyddsmyndigheten DFS 2004:4) in a protocol approved by the local Ethical Committee (Umeå djurförsöksetiska nämnd, Permit number A12-13, A80-14 and A79-14.

### Statistical analyses

Statistical analysis was conducted using Graphpad Prism 6 software and *P* values less than 0.5 were considered significant. All two-group comparisons in *Candida* CFU data, inflammatory score, ALT data, and ELISA data were conducted using the unpaired, two-tailed student’s *t*-test. Comparisons of WT and *S100A9*^*-/-*^ mice data were performed using One-way ANOVA multiple comparisons analysis as specified in figure legends. In all comparisons, the sample size is specified in figure legends, and a *P*<0.05 was considered significant. *p<0.05; **p<0.01; ***p<0.001; ****p<0.0001.

### Yeast strains and growth conditions

*C. albicans* clinical isolate strain - SC5314 was cultured overnight in YPD (1% yeast extract, 2% bacto-peptone and 2% glucose) at 30°C. The *Candida* cells were washed three times in PBS prior use in all assays. Cell numbers were calculated using Vi-CELL Cell Viability Analyzer (Beckman Coulter AB).

### Animal infections and isolation of bone marrow-derived macrophages, and tissue analyses

All mice were maintained according to a previous report (23) at Umeå Centre for Comparative Biology (UCCB), Umeå University, Umeå, Sweden. If not otherwise stated mice were infected intraperitoneally with 3×10^6^ *C. albicans* cells per g mouse from an overnight culture in YPD. For intravenous infection (only Fig. S4) mice were challenged intravenously with 2.5 × 10^3^ *C. albicans* cells per g mouse.

Primary macrophages (BMDMs) cells and differentiated as described in a previous report (24). BMDMs flow cell cytometry (FACs) was conducted using BD LSR II flow cytometer (BD Biosciences, San Jose, CA), using propidium iodide (24), on paquinimod treated cells at indicated concentrations.

For tissue damage and fungal load, analyses were conducted according to a previous report (25), with 3×10^6^ *C. albicans* cells intraperitoneal injection per g of mouse after 24 hours of infection (26). Blood ALT levels were measured using vetscan VS2^TM^ (SCIL animal care company) as previously reported (27–29).

Histological preparations and inflammatory score analyses were conducted as in previous reports (15,30). For the inflammatory score, Whole sections were analyzed for inflammation and scored under the supervision of a specialized animal pathologist. The sections from each animal were scored as zero if they had no inflammatory cells present in the tissue, one for a few inflammatory cells (1–20 cells), two for moderate cell infiltration (21–40 cells), three for a large number of inflammatory cells (41–60 cells), and four if inflammation was spread all over in the tissue (61 cells) (30).

### Cytokine and chemokine quantification

BMDMs were seeded at 1×10^5^ cells per well in 96-well microplates and then infected with *C. albicans* at an MOI of 1 for 24 hours. Cell-free culture supernatants were harvested after *C. albicans* infection. Supernatants were analyzed for indicated cytokines of chemokines by ELISA (Biolegend-ELISA MAX™ 9727 Pacific Heights Blvd, San Diego, CA, USA), or Pro-Mouse cytokine BioPlex^®^ 200 multiplex (Bio-Rad Laboratories) according to the manufacturer’s instructions.

### Generation of recombinant S100A8

The S100A8 and S100A9 gene encoding for monomers of the mouse dimer S100A8/A9 (UniProt P27005, P31725) was synthesized, codon harmonized and purchased from DNA2.0. The gene were GST tagged and cloned into *E. coli* BL21 strain. The cloned cells were auto-induced overnight. The cells were pelleted and suspended in 10 mg/ml of 1×PBS supplemented with DNase and protease inhibitors. The cells were lysed on ice by sonication (Branson Digital sonifier; 10 mm horn, 50% power) for 6 min with alternating 10 s pulses and pauses. The lysate was clarified by centrifugation at 23,000g for 20 min at 4°C. The clarified lysate was filtered through a 0.45 µm syringe filter and batch bound to 2.0 ml GST sepharose (∼25mg protein/ml) for 2h at 4°C gravity flow over column and washed with 30 column volumes of PBS. Samples were eluted in 4-6 ml fractions with 50 mM Tris pH 8 and 10 mM glutathione. For each fraction, the resin was incubated with elution buffer for 10 min prior to collecting the flow through. The fractions were analyzed at A280. The pooled fractions of protein were cleaved (1:100) using protease 3c (PreScission®protease) to remove the GST tag and the fractions of GST tag free protein were pooled. The protein was then separated by size-exclusion chromatography (SEC) using a Superdex 75 16/600 column (GE Healthcare Life Sciences, UK) equilibrated with PBS pH 7, at a flow rate of 0.5 ml/min. The fractions containing purified S100A8 (rA8) and S100A9 (rA9) were concentrated using an Amicon Ultra-4 centrifugal filter device with a 3 kDa molecular-weight cutoff (Millipore). The primary sequence, the intact mass and the presence of product were confirmed by mass spectrometry using an ABI 4800 MALDI tandem time-of-flight mass spectrometer. Recombinant proteins were screened for endotoxin contamination and levels were below 0.6 pg per µg protein, as previously recommended (31).

## Results

### A disseminated fungal peritonitis model to determine roles of S100A8/A9

During IAC, fungal cells dissemination reach the liver and other organs via lymphatics or bloodstream (6). Liver tissue damage and leukocytosis are hallmarks of deep-seated and systemic *C. albicans* infection (32). To establish a fungal IAC model and determine the role of S100A8/A9 in systemic inflammation, we used *S100A9*-deficient mice (*S100A9*^*-/-*^) that also lack the S100A8 protein on the protein level despite normal S100A8 RNA level and thus actually represent a functional S100A8/A9 double knockout mouse strain (33). Normally expressed S100A8 protein in S100A9^-/-^ mice is rapidly degraded in the proteasome, avoiding systemic overwhelming immune responses. However, under chronic TNF conditions, this degradation process is insufficient leading to severe phenotypes in artificial mouse strains (TTP^-/-^/S100A9^-/-^ or itghTNF/S100A9^-/-^) indicating that heterodimer activity must be regulated tightly to restrict operating range (13).

Using wildtype (WT) and *S100A9*^*-/-*^ mice, *C. albicans* intraperitoneal injections were administered, liver tissues collected after 24 hours of infection and indicators of sepsis were measured. The infection was determined to be systemic using phenotypic analysis of mice and histopathological analysis from hematoxylin-eosin stained (H & E stain) liver sections after 24 hours. The development of eye exudates was an indication of fungal dissemination from the peritoneal cavity to other loci in the mouse (Fig. 1A). WT liver sections indicated higher numbers of leukocyte infiltration zones compared to *S100A9*^*-/-*^ sections (Fig. 1A, top panel). Conversely, there was visual evidence of more fungal cells in the *S100A9*^*-/-*^ livers compared to WT (Fig. 1A, bottom panel). The inflammatory score (Fig. 1B) showed a large number of inflammatory cell infiltrates (level 3) in WT *C. albicans* infected samples compared to lack of inflammation (level 1) observed in *S100A9*^*-/-*^ mice.

**Figure 1.**
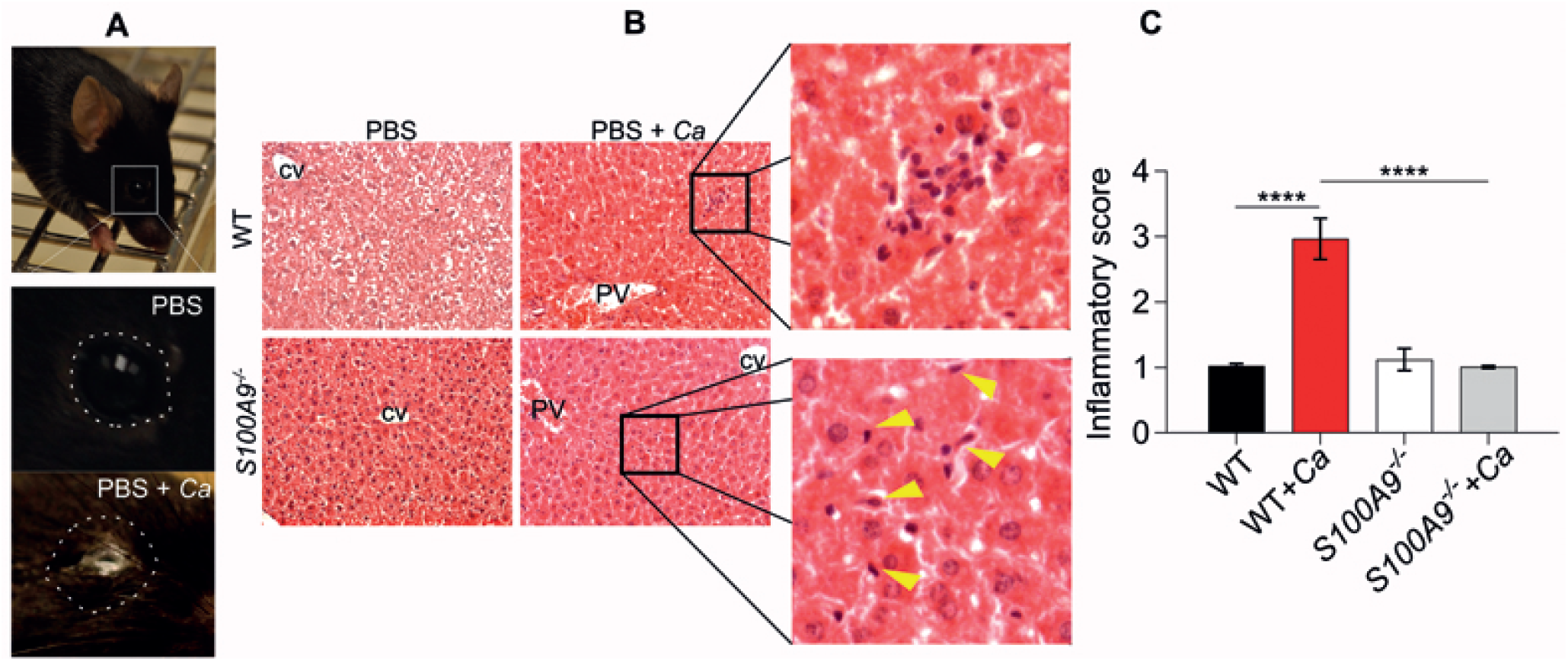
Disseminated *C. albicans* peritonitis mouse model depicting the necessity of S100A8/A9 in inflammation. Intraperitoneal (IP) *C. albicans* infection (3 × 10^6^ cells per g mouse) in WT or S100A8/A9-deficient mice after 24 hours. **A)** Fungal eye exudate as evidence for peritoneal cavity failure to contain C. albicans infection after 24. Shown is the clear eye of uninfected (PBS) compared to the white pus in infected wild type (WT) mouse. **B)** Representative photomicrographs of hematoxylin-eosin (H&E) stained liver sections of uninfected and infected wildtype and s100A9 mutant used in quantifications in **C)**. Zoomed images show one of many observed increased inflammatory cell infiltrates in WT infected cells (top panel), and arrows indicate yeast or hyphal cells (bottom panel) at 20x magnification. CV: central vein, PV: portal vein. **C)** An inflammatory score of H&E-stained liver sections of uninfected and infected WT and S100A9-/- mutant mice. A score of 2 = moderate cell infiltration, >3 = large number of infiltrates, 4 = full tissue inflammatory infiltration. Data from 10x microscopy image fields of 2 sections from mice n=2 (WT and *S100A9* ^*-/-*^) and n=6 (WT + Ca and *S100A9* ^*-/-*^ + Ca). Error bars indicate standard deviation and ****p< 0.0001

Blood alanine aminotransferase (ALT) levels in the blood are an indicator of liver damage (1,29). *C. albicans* infected WT mice (101.6U/L) showed elevated levels compared to *S100A9*^*-/-*^ mice (37.2U/L) which showed similar ALT levels to uninfected mice, suggesting lack of systemic tissue damage in the S100A8/A9-deficient animals (Fig. 2A). The ability of the host to clear the infection or organ microbial load is related to the colony-forming units (CFUs) (34). *S100A9*^*-/-*^ mice had a significantly higher fungal load (2.5 fold of average levels) compared to WT mice (Fig. 2B).

**Figure 2.**
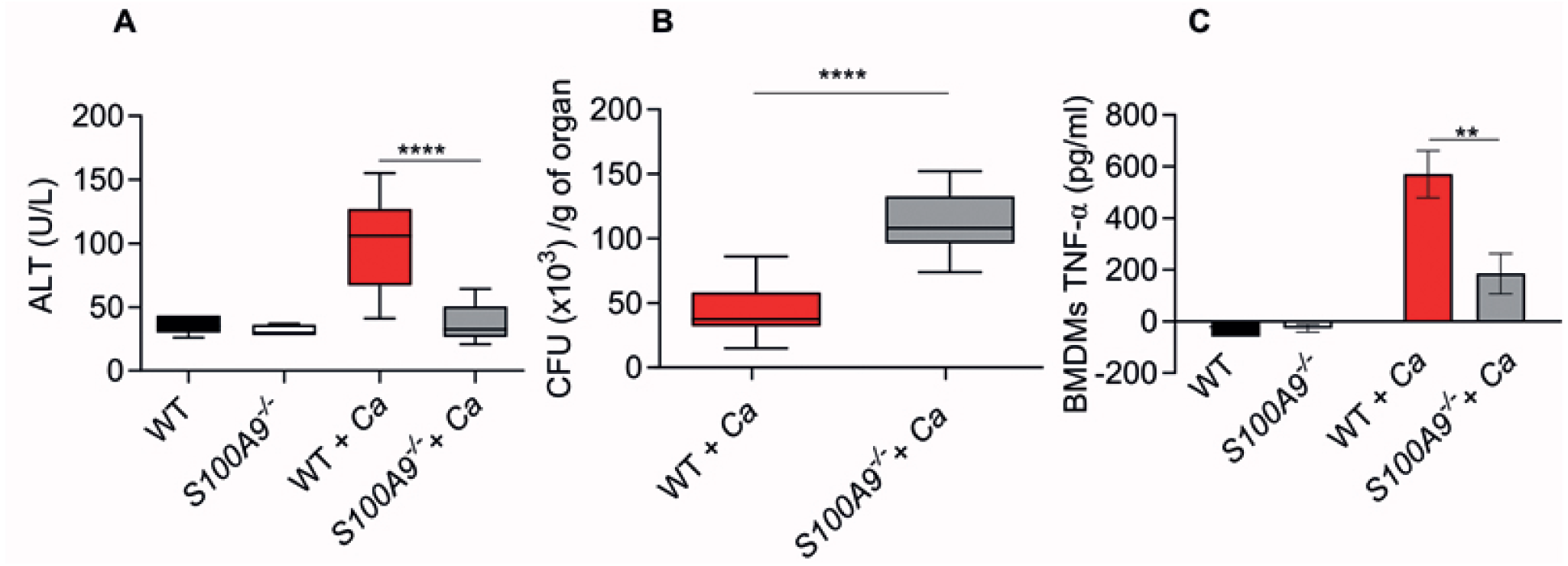
S100A8/A9 is required for inflammatory induced collateral tissue damage (CTD) and fungal clearance in the liver. **A-B)** WT and S100A9-/- mice were infected with 3 × 10^6^ cells per g of mice. A) Alanine transferase levels (ALTs) used as an indicator of liver tissue damage as a result of *C. albicans* infection. Plasma ALT levels were analyzed from 100µl of plasma in a vetscan rotor 24 h post-infection. **B)** Organ levels of colony-forming units (CFUs) as a measure of host fungal burden. *C. albicans* CFUs in homogenized infected livers of indicated mice were quantified. **A-B)** Box- and whisker plots show the smallest observation, lower quartile, mean, upper quartile and largest observation. n = 7 (WT and S100A9-/-), and n= 10 (WT and S100A9-/- + Ca) total mice. **** p-value =<0.0001. **C)** *C. albicans in vitro* infection of immune cells induces the production of less pro-inflammatory cytokines. WT and S100A9-/- bone marrow-derived macrophages (BMDMs) were infected with *C. albicans* (MOI 1). TNFα levels were measured from supernatants at 24-hour post-infection. ** p-value =<0.01, n = 3.

Macrophages are sentinels for immune signaling that leads to leukocytosis but may cause problems when uncontrolled (35). The pro-inflammatory cytokine response of WT and *S100A9*^*-/-*^ was determined using bone marrow-derived macrophages (BMDMs) to monitor the ability of primary macrophages to induce TNFα (Fig. 2C). Higher levels of TNFα (3.1 fold of average levels measured) upon *C. albicans* infection were induced by WT BMDMs (570 pg/ml) compared to *S100A9*^*-/-*^ BMDMs (185 pg/ml), suggesting a defect in TNFα induction in cells lacking S100A8/A9.

### S100A8/A9 enhances pro-inflammatory cytokine release of macrophages

The implications of the use of S100A8/A9 in recombinant protein therapy is unknown. To determine whether the S100A8 activity would aid the *S100A9*^*-/-*^ mutant in eliciting an appropriate cytokine response *in vitro*, BMDMs derived from S100A9-/- mice were infected with *C. albicans* and treated with rS100A8. Murine *S100A8* was expressed in *E. coli*, purified (Fig. S1) and verified by mass spectrometry to obtain functionally active homodimers as a substitute for the heterodimer as previously reported (36). We included analyses for cytokines typically released during early inflammatory responses (Fig. 3A-F).

**Figure 3.**
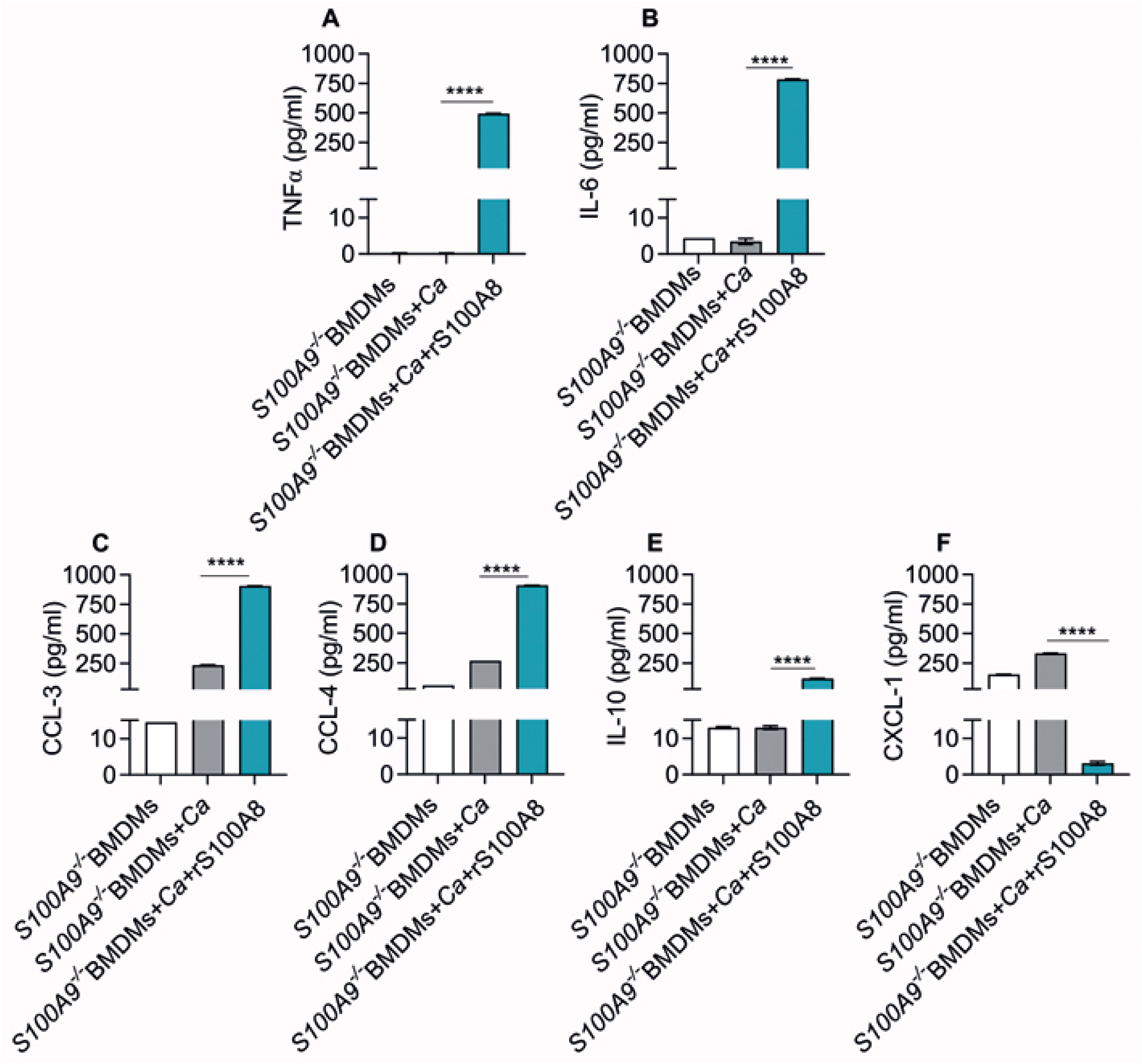
Recombinant S100A8 protein restores pro-inflammatory cytokine release of *S100A9*^*-/-*^ bone marrow-derived macrophages upon *in vitro C. albicans* infection. Cytokine levels **A)** TNFα, **B)** LI-6, **C)** CCL3, **D)** CCL-4, **E)** IL-10, and **F)** CXCL-1 in supernatants of bone marrow-derived macrophages (BMDMs) infected with *C. albicans* (MOI 1) were measured 24-hour post-infection. A Pro-Mouse cytokine BioPlex^®^ 200 multiplex array (Bio-Rad, Hercules, CA) was used to detect and quantify *S100A9*^*-/-*^ mouse cytokines infected with *C. albicans* with and without additional treatment with rS100A8 (10 µg/ml). **A-F)** n=3, \**p*-value =<0.05 \*\*\*\**p*-value =<0.0001.

Considerable increase of TNFα levels was obtained in *S100A9*^*-/-*^ BMDMs treated with rS100A8 (Fig. 3A) comparable to WT levels upon *C. albicans* infection (Fig. 2C), and IL-6 was also strongly enhanced (Fig. 3B). In addition, *S100A9*^*-/-*^ BMDMs infected with *C. albicans* and treated with rS100A8 released higher levels of chemokines *MIP1α* (*CCL3*) and *MIP1β* (*CCL4*), crucial for the recruitment of various leukocyte subpopulations (Fig. 3C and 3D) as compared to infected and untreated control macrophages. Also monitored was the induction of the anti-inflammatory cytokine IL10, which showed low but significant levels induced when mice were treated with rS100A8, while CXCl-1 levels declined (Fig. 3E and 3F). Induction of pro-inflammatory cytokines and modulation of tissue damage by rS100A8 suggest that S100A8/A9 heterodimers are main contributors of the acute pro-inflammatory response during *C. albicans* infection.

### Paquinimod reduces cytokine release of C. albicans-infected macrophages

To potentially reduce ICTD during IAC pharmacological intervention blocking the pro-inflammatory activity of S100A8/A9 could be used. Hence. the activity of paquinimod, a novel anti-inflammatory compound initially developed against Systemic Lupus Erythematosus (SLE) that targets S100A9, one subunit of the S100A8/A9 complex, in liver, lung, heart and skin (37), was tested *in vitro*. The percentage of dead cells (propidium iodide positive cells) using flow cytometry analysis (FACS) was assessed on treatment of WT BMDMs with various drug concentrations of paquinimod. Among the concentrations used, no significant toxic effects were observed (Fig. S2). However, there was significant TNFα reduction at 300 µg/ml and 930 µg/ml paquinimod concentrations compared to untreated *C. albicans* infected BMDMs (Fig. 4A). CCL-3 and IL-10 were also reduced upon addition of 930 µg/ml paquinimod confirming the activity for other cytokines and chemokines. Thus, paquinimod is likely able to block S100A8/A9-mediated cytokine release during *C. albicans* infection.

**Figure 4.**
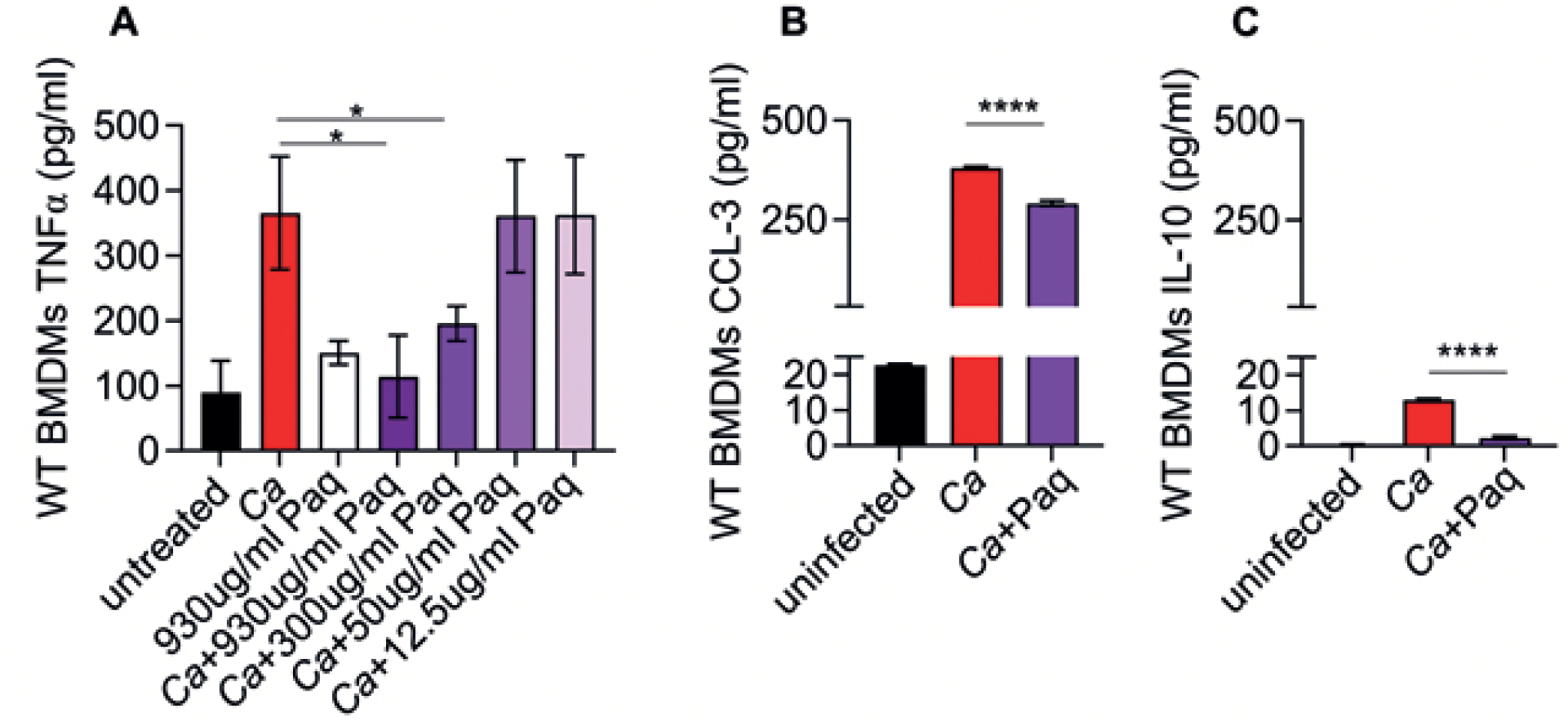
Quinoline-3-carboxamide paquinimod (Paq) reduces pro-inflammatory cytokine release of WT bone marrow-derived macrophages upon *in vitro C. albicans* infection. **A)** WT BMDM were infected and treated with different Paq concentrations (930µg/ml – 12.5 µg/ml). TNFα was measured from supernatants 24 h post-infection. **B-C)** WT BMDM were infected and treated with 930 µg/ml or 0 µg/ml Paq and release of B) CCL-3 and C) IL-10 was measured from supernatants after 24 h. **A-C)** n=3. \**p*-value =<0.05, \*\*\*\**p*-value =<0.0001.

### S100A8/A9 is responsible for ICTD during disseminated peritonitis

The active role of S100A8/A9 as alarmin during IAC is currently unknown. To test the usefulness of recombinant protein therapy, we determined the effects of recombinant S100A8 protein (rS100A8) on disseminated *C. albicans* infection. Purified rS100A8 protein (100μl of a 100 μg/ml solution) was injected into *C. albicans* infected WT and *S100A9*^*-/-*^ mice. Treatment of *S100A9*^*-/-*^ mice with rS100A8 showed a higher level of ALT (5.9 fold of average levels measured) compared to untreated with levels not significantly different to WT suggesting that constitutively active S100A8 homodimers mimic heterodimers to cause WT levels of liver damage (Fig. 5A) and that the S100A8/A9-mediated effect on ICTD is direct rather than indirect. S100A8/A9 is an antimicrobial protein against *C. albicans* (15). In this context, the fungal clearance defect observed in *C. albicans* infected *S100A9*^*-/-*^ mice was remedied, reducing fungal load after treatment with rS100A8 (Fig. 5B).

**Figure 5.**
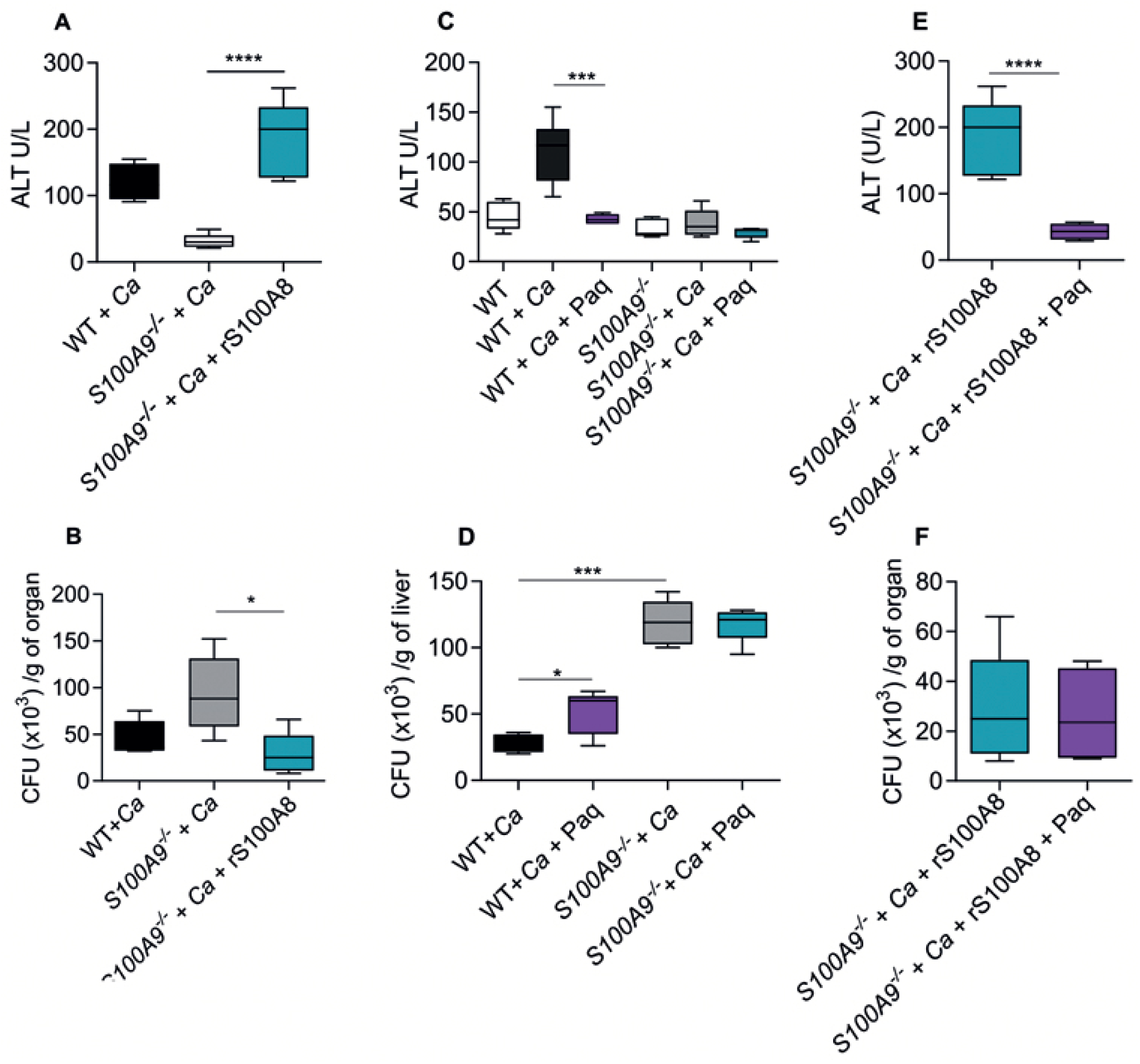
Recombinant S100A8 and paquinimod modify collateral tissue damage in experimental disseminated *C. albicans* peritonitis. **A).** Effect of recombinant soluble rS100A8 on *C. albicans* induced collateral tissue damage. Protein was intraperitoneally administered (approximately 10 µg per mouse) to WT and *S100A9*^*-/-*^ mice intraperitoneally infected with *C. albicans* (3 × 10^6^ cells per g mouse) and shown are plasma ALT levels. **B)** Effect of rS100A8 on host fungal clearance. Shown are CFUs from homogenized infected livers of indicated mice **A-B**) Data are presented as a box- and whisker plots showing the smallest observation, lower quartile, mean, upper quartile and largest observation, statistical significance was analyzed by one-way ANOVA n = 5 mice per group in 3 separate experiments, **** *p*-value =<0.0001. **C-D)** Effects of paquinimod (Paq) on *C. albicans*-induced collateral tissue damage and fungal clearance in intraperitoneally infected mice. Paq was administered intraperitoneally (30mg/kg). Shown are plasma ALT levels **C)** and CFUs from homogenized infected livers **D)** from *C. albicans* infected WT and *S100A9*^*-/-*^ mice. E-F) Effect of Paq on recombinant S100A8/A9 induced collateral tissue damage during *C. albicans* infection. Note that **E)** contains *S100A9*^*-/-*^ controls initially presented in **A)** as experiments were conducted together. Shown are plasma ALT levels **E)** and CFUs from homogenized infected livers **F)** from *C. albicans* infected *S100A9*^*-/-*^ mice treated with rS100A8 (10µg per mouse) with our without Paq treatment. Data are presented as a box- and whisker plots showing the smallest observation, lower quartile, mean, upper quartile and largest observation, statistical significance was analyzed by student t-test n = 5 mice per group, \**p*-value =<0.05, **** *p-*value =<0.0001.

### Paquinimod therapy reduces ICTD induced by disseminated peritonitis

Targeted deletion of *S100A9* improves survival in mouse models of bacterial-induced sepsis (11,38). Higher lethality of a disseminated infection is often due to the inability to contain inflammation-induced organ and tissue damage by the host (1). There are no specific immunotherapies against disseminated infections, such as sepsis, and often management focuses on containing the infection through source control and antibiotics or antifungals plus organ function support (39). We hypothesized that using an anti-inflammatory drug in WT mice may mimic the beneficial aspect (lack of tissue damage) observed in the *S100A9*^*-/-*^ mutant might extend host survival from disseminated IAC. Treatment of intraperitoneally infected mice with paquinimod led to complete elimination of ICTD as indicated by reduced ALT plasma levels in infected mice to levels of uninfected mice (Fig. 5C). As expected, paquinimod treatment did not have an effect on ALT levels in infected *S100A9*^*-/-*^ mice (Fig. 5C). In addition, paquinimod treatment affected fungal burden in the liver of infected WT and *S100A9*^*-/*-^ mice only moderately (Fig. 5D), probably due to reduced S100A8/A9-dependent activation of fungi-eliminating immune cells. Of note, the CFU count in livers of *S100A9*^*-/*-^ mice was significantly higher than in livers of WT mice, suggesting that ICTD after 24 h of infection is not strictly correlated to fungal burden, but depends in this setting to a significant proportion on S100A8/A9 activity.

Paquinimod and other quinoline-3-carboxamides have been described as specific binders of S100A9 (40). As rS100A9 does not form functional homodimers *in vitro* and hence could not lead to a reduction of ICTD in experimental *C. albicans* peritonitis (Fig. S3), we used rS100A8. Interestingly, the effect of rS100A8 injection on ALT plasma levels in infected *S100A9*^*-/*-^ mice could be reverted by treatment with paquinimod (Fig. 5E), but expectedly had no effect on fungal burden in the liver (Fig. 5F). It is possible that paquinimod binds recombinant mouse S100A8 with somewhat higher affinity than recombinant human S100A8 which possibly could explain the described effect.

### Paquinimod has a moderate effect on survival in disseminated peritonitis

To determine whether paquinimod could be used to alleviate sepsis derived from fungal peritonitis, infected wildtype mice were treated with paquinimod every 24 hours with paquinimod (Fig. 6A). The treatment resulted in a moderately increased survival rate after five days post-infection with a statistically significant effect using log-rank test (Fig. 6B). Although the compound did not prevent all mice from succumbing to systemic infection, between 24 and 48 hours treated mice survived longer compared to untreated mice (Fig. 6B). Despite the similar weight loss in infected mice observed between treated and untreated groups (Fig. 6C), treated mice were generally more active during *C. albicans* infection. Hence, our data suggest that anti-S100A8/A9 therapy could be a useful strategy to increase the therapeutic window for antifungal treatment during IAC, but probably could not serve as a standalone therapeutic approach.

**Figure 6.**
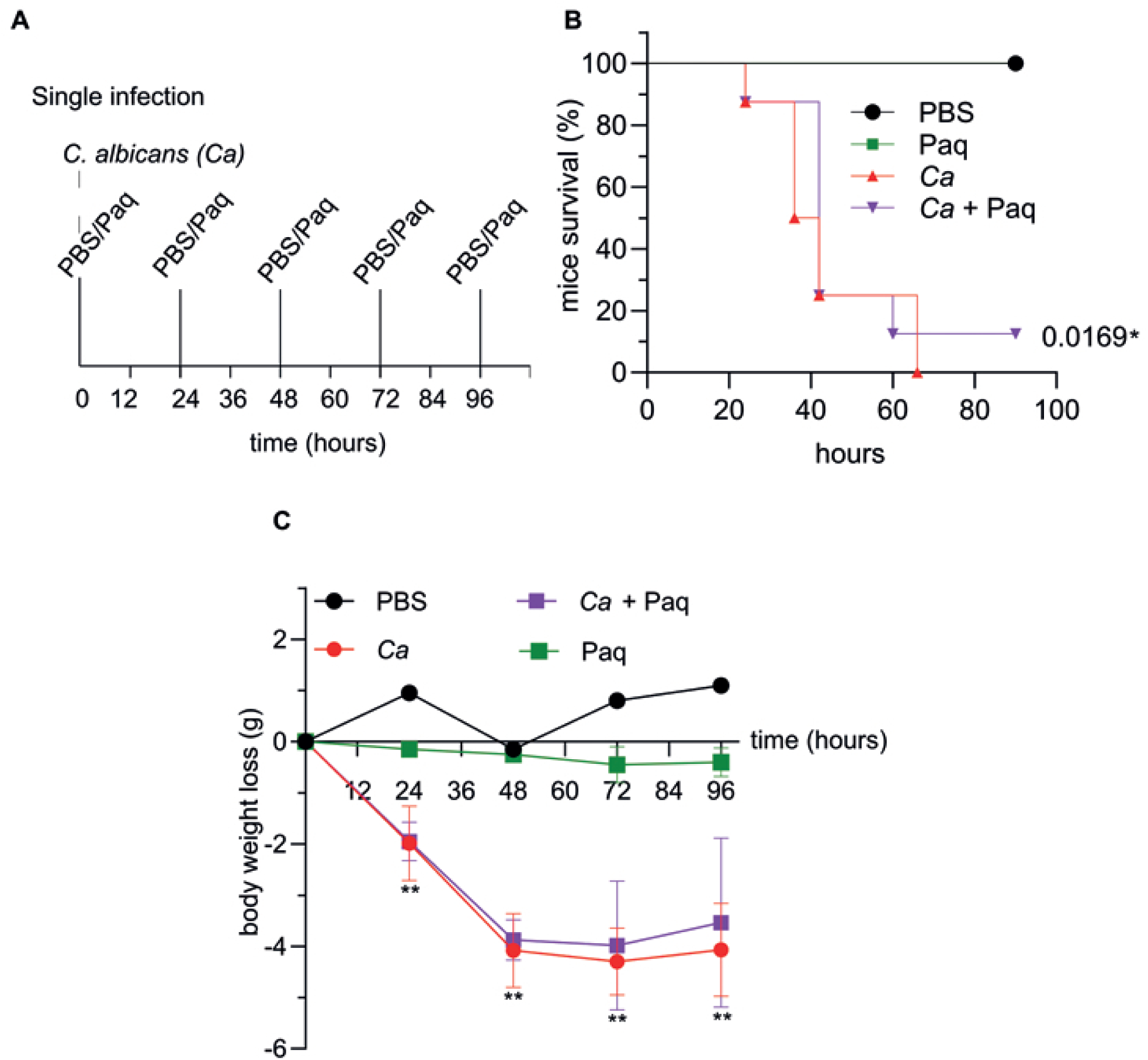
Inhibition of S100A8/A9 by paquinimod moderately increases host survival from disseminated *C. albicans* peritonitis. **A-D)** WT mice infected with *C. albicans* were treated with Paq (30mg/kg) at 24-hour intervals up to 5 days, and mice weight and survival were monitored. **A)** Paq mice treatment strategy. **B** Survival B) and sequential body weight C) of WT mice infected and treated at the indicated timeline. Survival is moderately but significantly increased. n = 8 mice per group. For survival log-rank test *p-*value = 0.0169, for mouse weight grams lost over time is shown. **p-value <0.01. Shown are mean and SD.

## Discussion

This work characterizes the role of immune-modulating S100A8/A9 on host resolution of inflammation from a peritoneal-derived disseminated *C. albicans* infection. The gastrointestinal commensal nature of *C. albicans* requires that mucosal damage and neutropenia are achieved for *C. albicans* dissemination (41). Most experimental models studying systemic fungal diseases use intravenous (IV) injection of fungal cells which bypass mucosal host defenses and establish an infection predominantly in the kidney and in the brain (42). This study utilized an IAC model to induce systemic inflammation mimicking a severe clinical concern of postoperative *Candida* peritonitis (43). Similar to IV, IP-induced infection allows rapid blood dissemination of pathogens with exposure to an active population of phagocytes, complement cascade and the potential for abscess formation in the peritoneal cavity (43). The peritonitis infection model presented here aimed to breach the immune barriers in the peritoneal cavity, and this was phenotypically clear in the dissemination of *C. albicans* to the liver and to the eyes of infected animals (Fig. 1 and 2). The IP injection route allows pathogen exposure to active phagocyte populations in the peritoneum and the potential for the host to contain the infection through the formation of abscesses (6). In intravenous infection, in contrast to the IP injection route, ALT plasma level did not increase above uninfected control within the first 48 h of infection (Fig. S4) and hence the peritonitis infection model is suitable to describe acute and severe onset of disseminated infection leading to ICTD.

WT liver fungal clearance was ∼3 times more efficient when compared to the S100A8/A9-deficient mice (Fig. 2B). The various leucocyte infiltration zones observed in WT liver tissues compared to *S100A9*^*-/-*^ mice (Fig. 1A) indicated that the recruitment of higher levels of leukocytes coincides with fungal clearance in WT mice. These findings suggested that the presence of S100A8/A9 is required for fungal clearance during disseminated infection, and supports our previous findings that implicated antimicrobial properties of S100A8/A9 during candidiasis (15). Notably, presence of S100A8/A9 in WT mice resulted to significantly more liver damage compared to *S100A9*^*-/-*^ mice as indicated by ALT plasma levels despite higher fungal load in the liver (Fig. 2A and B), implicating that the induction of S100A8/A9 inflammatory responses modulates processes which contribute to tissue damage. These findings support the association of the S100 family of proteins with inflammatory disorders (13), and our data indicate that S100A8/A9 activity depends on modulation of both antimicrobial and inflammatory activity. These insights would especially be critical in the case of immunocompromised patients.

The IP treatment with rS100A8 of *C. albicans-*infected *S100A9*^*-/-*^ mice resulted in fungal clearance similar to WT mice. Since rS100A8 showed significant *in vivo* activity, we used *S100A9*^*-/-*^ bone marrow-derived macrophages in a *C. albicans ex vivo* infection assay, and showed that rS100A8 mediates macrophage activation as indicated by the higher levels of the chemokines CCL-3 and CCL-4, pro-inflammatory cytokines IL-6 and TNFα, as well as the anti-inflammatory cytokine IL10 (Fig. 3). The induction of IL10 by rS100A8 is consistent with homeostatic management of systemic inflammation (44). Balancing of pro- and anti-inflammatory cytokines is a common theme of host factors with local and system-wide effects as exemplified by interferon signaling pathways during viral infections (45). Albeit administration of rS100A8 led to enhanced fungal clearance in the liver of infected S100A8/A9-deficient mice, the activity of S100A8 has the potential to induce dire effects on host-mediated tissue damage which probably do not warrant recombinant therapy with rS100A8 in immunocompromised individuals.

Our data demonstrate that ICTD, which was alleviated in S100A8/A9-deficient mice, could be fully reintroduced by injection of recombinant protein (rS100A8). Thus, the reported effect on ICTD in an experimental model of IAC is, at least to a large proportion, dependent on S100A8/A9 and pharmacological targeting of the protein complex could be beneficial in patients with IAC. To demonstrate this, we used paquinimod; a compound applied for treatment of chronic inflammatory diseases (17,20). As paquinimod is a specific binder for S100A9 (40), it blocks the pro-inflammatory activity of the S100A8/A9 complex *in vivo*. Surprisingly, we found that paquinimod also reverted ICTD in infected S100A9-/- mice previously treated with rS100A8 (Fig. 5E). Bjork *et al*., showed strong binding of paquinimod to human and mouse S100A9 and negligible binding to human S100A8 *in vitro*; however, binding to mouse S100A8 was not investigated (40). Notably, it may also be possible that paquinimod exerts other immune-modulatory effects that give rise to the reduction seen in Fig. 5E. No toxicity was observed against bone marrow-derived macrophages (BMDMs) at the concentration used in animal infection experiments and paquinimod activity affected the inflammatory modulators as observed in the reduction of CCL-3 and IL-10. No adverse effects on fungal load in livers of infected animals were seen either (Fig. 5D). Hence, paquinimod represents a potential adjuvant therapeutic option that dampens the host responses to avoid tissue damage and increases the therapeutic window (Fig. 6B) for targeting the pathogen with antifungal drugs, in particular, since the compound had no adverse effect on fungal clearance.

This study showed the dilemma of the protective and harmful roles of S100A8/A9 in *Candida albicans*-induced fungal peritonitis (Fig. 6). Fungal peritonitis that is not contained by host defenses leads to pathogen infiltration of host organs (Fig. 7.1). Fungal-host interaction induces the systemic release of S100A8/A9 (15) for fungal clearance and ALTs into the systemic circulation (Fig. 7.2-7.3). Local innate immune cells (represented in this study as primary macrophages respond with a cytokine ‘storm’ of pro-inflammatory cytokines (Fig. 7.4-7.5), that leads to increased leukocyte infiltrates systemically (Fig. 7.6), and promote increased tissue damage in the effort of fungal clearance (Fig. 7.7). Hence, our study presents both the potential for a systemically functional protein, evidence for potential antimicrobial recombinant protein therapy, and adjuvant treatment as paquinimod-mediated inhibition of S100A8/A9 showed reduction of ICTD (Fig. 7). Considering the high mortality (>50%) due to do fungal sepsis (1), future adjuvant therapies similar to paquinimod could be the key against peritonitis and other severe inflammatory malignancies.

**Figure 7.**
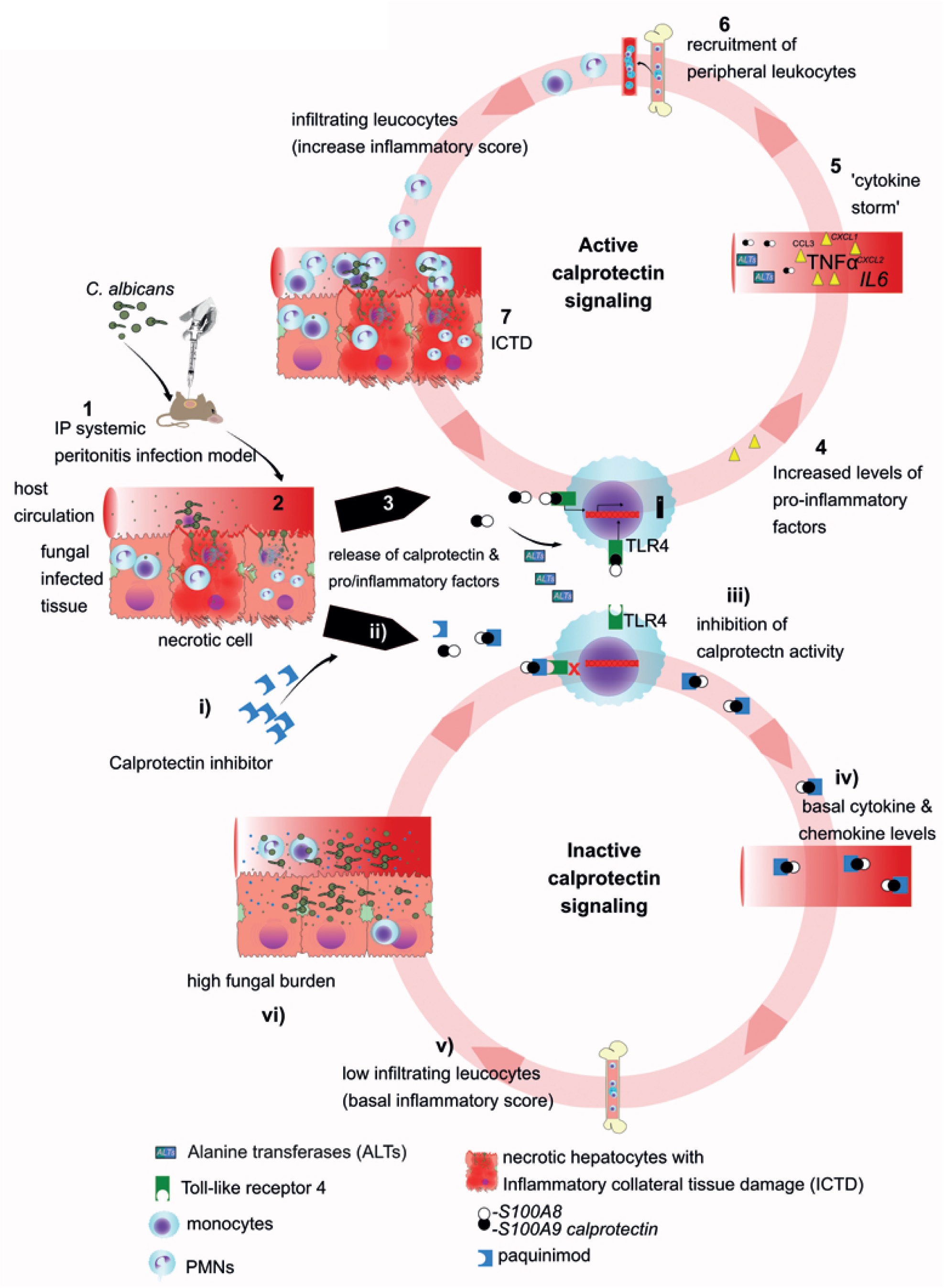
Study summary. Using a systemic fungal peritonitis mouse model **(1)**, findings implicate S100A8/A9 as a systemically essential protein **(2-6, i-vi))** for the control of cytokine and chemokine modulation **(4-5)**, organ leukocyte recruitment **(6)**, host tissue damage **(7)** and fungal clearance **(vi)**. In systemic intra-abdominal originating candidiasis, the cost of fungal clearance (antimicrobial/active S100A8/A9 signaling) a feedback-loop that increases tissue damage, while the cost of decreased tissue damage (anti-inflammatory/ inactive S100A8/A9 signaling) is a higher fungal burden (Fig. 1-5). Both are lethal if not controlled (Fig. 5B). Future studies should focus on fine-tuning recombinant therapy in cases where active S100A8/A9 signaling is inadequate, and anti-inflammatory treatments like paquinimod, in conjunction with antifungal treatments to increase host survival time to aid treatment.

## Acknowledgments

The authors thank José Pedro Lopes and Sujan Yellagunda for support with animal experiments. We acknowledge essential advice and the kind gift of paquinimod from Active Biotech. We acknowledge support for heterologous protein expression of the Protein Expertise Platform (PEP) at Umeå University. Furthermore, we are grateful for funding provided to CFU by the Swedish research council VR-M 2014-02281 and VR-M 2017-01681, the Kempe Foundation SMK-1453, the Åke Wiberg Foundation M14-0076 and M15-0108, and the Medical Faculty of Umeå University 316-886-10. NU and MS held a postdoctoral fellowship received in competition from UCMR. The funders had no rule in study design nor analysis and interpretation of the results.

## Author Contribution

CFU provided funding and together with MS and NU designed the study, MS and NU collected and analyzed the data. TV and JR contributed to the conceptualization of the study. Manuscript drafting by NU, MS and CFU. SH contributed preparation of histological sections. MJN provided data from intravenous infection. All authors contributed to data analysis, drafting and critically revising the paper, gave final approval of the version to be published, and agree to be accountable for all aspects of the work.

