## Supplementary Figure S1 for "S100A8/A9 modulates inflammatory collateral tissue damage during intraperitoneal origin systemic candidiasis"

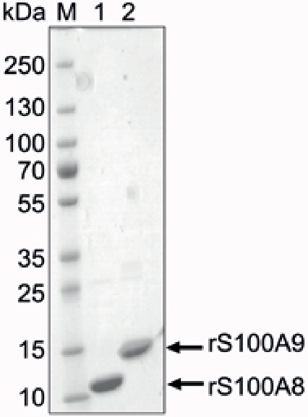


**Figure S1: Purity of recombinant calprotectin monomeric proteins rS100A9 and rS100A8.**

Coomassie-stained SDS protein gel with 5 µl purified mouse recombinant proteins rS100A8 (180 µg/ml) and rS100A9(490 µg/ml).
