## Supplementary Figure S2 for "S100A8/A9 modulates inflammatory collateral tissue damage during intraperitoneal origin systemic candidiasis"

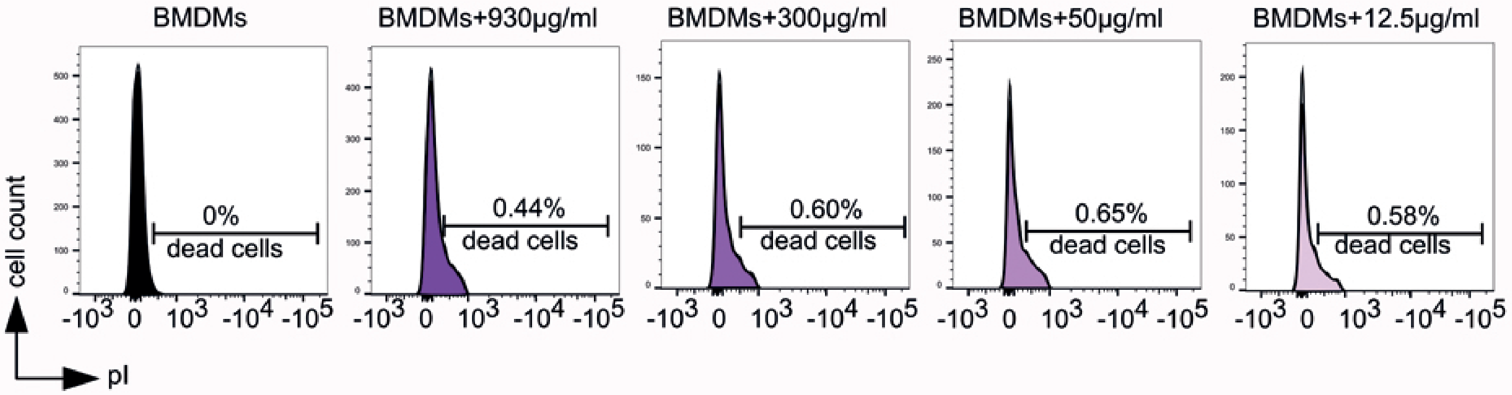


**Figure S2. Paquinimod at tested concentrations was not cytotoxic.**

Bone marrow-derived macrophages (BMDMs) were treated with indicated concentrations of paquinimod (Paq), stained with propidium iodide (pI) and analyzed using FACs live cell analysis. Shown are representative histograms and percent cell death due to paquinimod.
