## Supplementary Figure S3 for "S100A8/A9 modulates inflammatory collateral tissue damage during intraperitoneal origin systemic candidiasis"

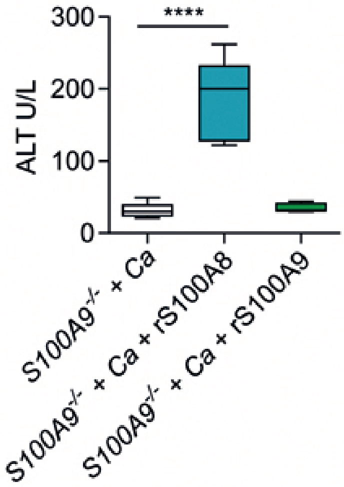


**Figure S3. Recombinant rS100A9 does not form functional dimers.**

Protein was intraperitoneally administered (approximately 10 µg per mouse) to WT and *S100A9*$^{-/-}$ mice intraperitoneally infected with *C. albicans* 3 X $10^6$ cells per g mouse and shown are plasma ALT levels. Control and rS100A8 treated sample are included in Fig. 5A, as experiments were conducted together. n = 5 mice per group in 3 separate experiments, **** *p*-value =<0.0001.
