## Supplementary Figure S4 for "S100A8/A9 modulates inflammatory collateral tissue damage during intraperitoneal origin systemic candidiasis"

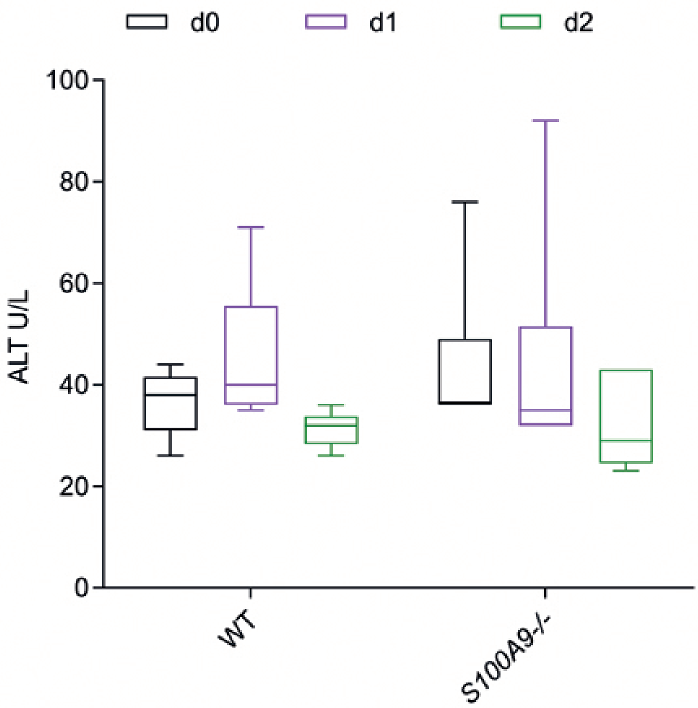


**Figure S4. ALT plasma level indicative for liver damage does not increase in the early phase of experimental systemic candidiasis (intravenous inoculation).**

WT and *S100A9*$^{-/-}$ mice were intravenously injected with 2.5 x $10^3$ *C. albicans* ells per g mouse. Plasma alanine transferase (ALT) levels were analyzed from 100µl of plasma in a vetscan rotor. Graph legend depicts d 0 = day 0, before infection; d 1 = 24 h post infection; d 2 = 48 h post-infection. n = 5 mice per group.
